# 3D Chromatin Dynamics during Innate and Adaptive Immune Memory Acquisition

**DOI:** 10.1101/2023.01.16.524322

**Authors:** Endi K. Santosa, Colleen M. Lau, Merve Sahin, Christina S. Leslie, Joseph C. Sun

## Abstract

Immune cells responding to pathogens undergo molecular changes that are intimately linked to genome organization. Recent work has demonstrated that natural killer (NK) and CD8^+^ T cells experience substantial transcriptomic and epigenetic rewiring during their differentiation from naïve to effector to memory cells. Whether these molecular adaptations are accompanied by changes in three-dimensional (3D) chromatin architecture is unknown. In this study, we combine histone profiling, ATAC-seq, RNA-seq and high-throughput chromatin capture (HiC) assay to investigate the dynamics of one-dimensional (1D) and 3D chromatin during the differentiation of innate and adaptive lymphocytes. To this end, we discovered a coordinated 1D and 3D epigenetic remodeling during innate immune memory differentiation, and demonstrate that effector CD8^+^ T cells adopt an NK-like architectural program that is maintained in memory cells. Altogether, our study reveals the dynamic nature of the 1D and 3D genome during the formation of innate and adaptive immunological memory.

## INTRODUCTION

Both innate and adaptive cytotoxic lymphocytes, namely NK cells and CD8^+^ T cells, respectively, are critical for host defense against pathogens. Although NK cells have traditionally been considered components of the innate immune system, accumulating evidence has demonstrated that these cells possess adaptive features, such as antigen-specificity, clonal expansion, and immunological memory across species^1,2^. More recent studies have suggested that the differentiation from naïve to effector to memory cells of innate and adaptive cytotoxic lymphocytes has parallel characteristics and is accompanied by significant transcriptomic and epigenetic transformation during viral infection^3–5^. Nevertheless, the molecular features of the dynamically regulated genomic regions, and which regions form networks to orchestrate optimal immune responses, remain unclear. Given that a variety of regulatory elements are critical for regulating gene expression and shaping the overall hierarchical structure of the 3D genome architecture, there is a need to understand how interactions between regulatory elements control the immune response in cell type- and cell state-specific manners.

Central to the 3D chromatin architecture are the interactions of regulatory regions that form chromatin loops, which in turn have the propensity to organize into topologically associating domains (TADs), and chromatin compartments that correspond to the transcriptionally active euchromatin and the transcriptionally inert heterochromatin^6^. Recent studies have shown that higher-order chromatin structures, which include chromatin compartments and TADs, undergo significant remodeling during development and lineage commitment^7,8^. However, whether similar processes occur in mature NK cells and CD8^+^ T cells that are experiencing extensive molecular adaptations that endow them with enhanced functionality is unknown. Here we perform an integrative analysis of transcriptome, chromatin accessibility, histone profiling, and 3D genome architecture to understand the entirety of immune memory differentiation through the chromatin lens in innate and adaptive cytotoxic lymphocytes.

## RESULTS

### Innate lymphocytes undergo coordinated histone remodeling during viral infection

To investigate the 1D epigenomic histone remodeling in innate lymphocytes during viral infection, we utilized CUT&RUN to generate longitudinal profiles of H3K27Ac (found at active promoters and enhancers) and H3K4me1 (putative enhancers) within Ly49H^+^ NK cells during mouse cytomegalovirus (MCMV) infection (Fig. 1A)^9,10^. Naïve, effector, and memory Ly49H^+^ NK cells were sorted (as shown in Supp. Fig. 1) and histone profiles showed expected peak distributions and high reproducibility across samples (Supp. Fig. 2A-B). To validate our findings, we first examined the profiles of H3K27Ac and H3K4me1 at the *Klrg1* locus, which is expressed only by a small fraction of naïve NK cells but is gradually upregulated in effector and memory Ly49H^+^ NK cells (Fig. 1B-C). We observed that the *Klrg1* locus and its surrounding loci were already highly decorated with H3K4me1 in naïve cells, and its occupancy was stable throughout the course of the infection, suggesting a permissive signature that may allow NK cells to readily upregulate this gene. Moreover, much like H3K4me1 occupancy, chromatin accessibility around the *Klrg1* locus also showed minimal changes. Unlike H3K4me1, we observed a highly dynamic H3K27Ac deposition that can be observed as early as day 2 post-infection (PI) and peaked on day 7 PI, with slightly delayed mRNA and protein expression early after infection that converged at later time points (Fig. 1B-C). In contrast, the coordination of H3K27Ac and the *Ifng* transcript occurred early after infection but diverged after day 2 PI. In this locus, H3K27Ac is still maintained at high levels in effector and memory cells despite the downregulation of *Ifng* expression (Supp. Fig. 2C). Altogether, these data highlight the complexity of gene regulation at these loci.

**Figure 1.**
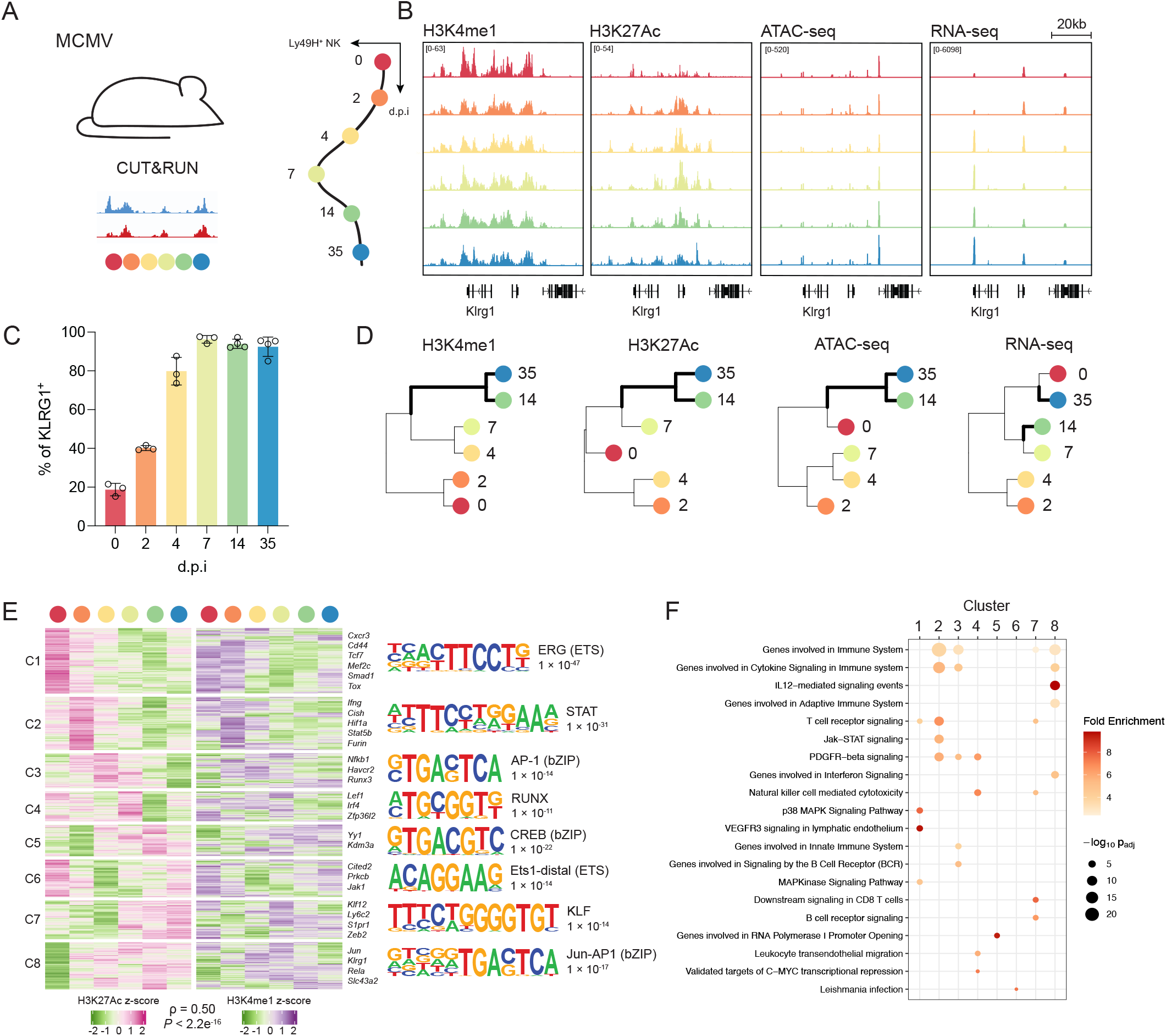
Innate lymphocytes undergo coordinated histone remodeling during viral infection. (A) Schematic of experimental setup and time points for MCMV time course (n = 2-4 samples per day for CUT&RUN). (B) H3K4me1, H3K27Ac, ATAC-seq, and RNA-seq tracks in the *Klrg1* and its surrounding loci of Ly49H^+^ NK cells at different time points following MCMV infection. (C) Percentage of KLRG1^+^ cells within naïve (d0) and antigen experienced (d2-35 PI) Ly49H^+^ NK cells during MCMV infection as assessed by flow cytometry. (D) Hierarchical clustering of H3K4me1, H3K27Ac, ATAC-seq, and RNA-seq based on normalized log_2_ counts. (E) Heatmap of H3K27Ac and H3K4me1 mean-centered normalized log_2_ counts (z-score) for high fold change regions (|log2FC| ≥0.5 and p_adj_ <0.05) based on H3K27Ac differential occupancy analysis (left). Top motif found in each cluster and its corresponding p-value (right). (F) Top 20 most enriched pathways across clusters shown as fold enrichment and −log_10_ adjusted hypergeometric p-value as calculated by Genomic Regions Enrichment of Annotation Tool (GREAT).

**Figure 2.**
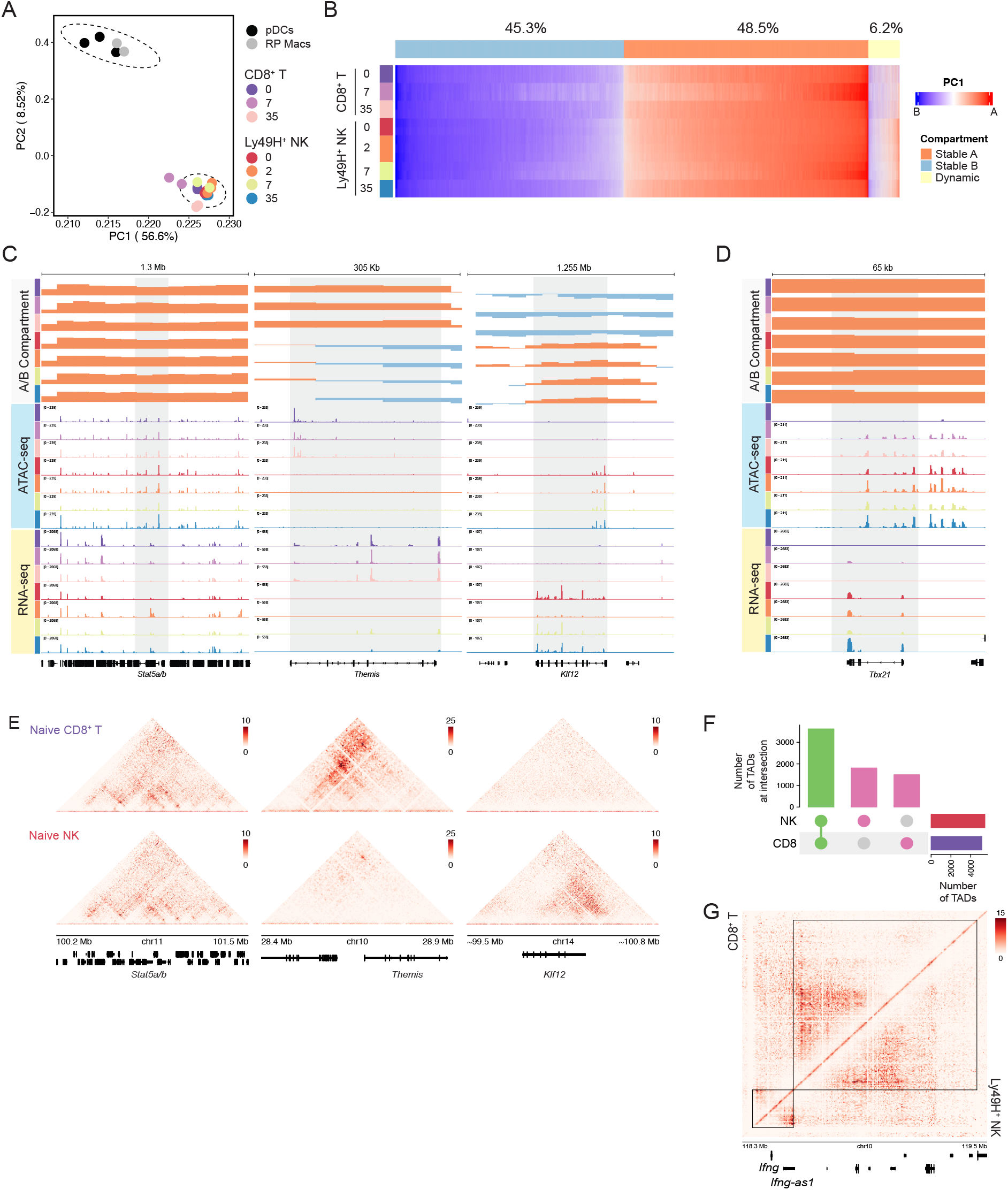
Higher-order 3D chromatin topology in innate and adaptive lymphocytes is stable during infection. (A) Principal Component Analysis of A/B compartment profiles of Ly49H^+^ NK cells, MCMV-specific CD8^+^ T cells, RP Macrophages, and pDCs. (B) Genome-wide A/B compartment heatmap of eigenvalues at 100kb bin. Eigenvalues were used to generate heatmap, and positive and negative values determine the direction of A or B compartments, respectively. Light blue, orange, and yellow bars represent ‘stable’ B, ‘stable’ A, and dynamic compartments, respectively. (C) Genomic tracks depicting A/B compartment, chromatin accessibility (ATAC-seq) and transcript (RNA-seq) on *Stat5a/b* (left), *Themis* (middle), and *Klf12* (right) loci of M45^H2-Db+^ CD8^+^ T cells (d0, 7, 35) and Ly49H^+^ NK cells (d0, 2, 7, 35) after infection. Orange bars represent A compartment, and light blue bars represent B compartment. (D) Genomic tracks depicting A/B compartment, chromatin accessibility (ATAC-seq) and transcript (RNA-seq) on *Tbx21* locus of CD8^+^ T cells (d0, 7, 35) and Ly49H^+^ NK cells (d0, 2, 7, 35) after infection. Orange bars represent A compartment, and light blue bars represent B compartment. (E) Contact heatmap matrix at 5kb resolution around *Stat5a/b, Themis*, and *Klf12* loci in naïve CD8^+^ T and naïve NK cells depicted as z-value. (F) Upset plot depicting shared and distinct high confidence TADs. Green bar shows TADs that are shared between NK cells and CD8^+^ T cells across time points, and pink bars represent counts of cell type-specific TADs throughout infection. (G) Contact heatmap matrix at 5kb resolution depicting shared TADs (black line) in naïve CD8^+^ T and naïve Ly49H^+^ NK cells around the *Ifng* and *Ifng-as1* loci depicted as z-value.

To investigate the dynamics of global epigenetic trajectories of these histone marks and chromatin accessibility throughout the course of MCMV infection, we performed principal component analyses (PCA) and unsupervised hierarchical clustering. Comparison of PCA between H3K4me1 and H3K27Ac showed similar trajectory patterns as naïve NK cells become effector and memory cells (Supp. Fig. 2D), reminiscent of the chromatin accessibility trajectory. Furthermore, global epigenetic commitment towards the memory fate (day 35 PI) also occurred as early as day 14 PI and was discordant with the transcriptome of differentiating NK cells (Fig. 1D). Given the similarity in the epigenetic trajectory patterns, we sought to understand the dynamic relationship between H3K27Ac and H3K4me1 during viral infection. To do so, we generated an atlas of overlapping peaks between H3K27Ac and H3K4me1 (Supp. Fig. 2E) and performed *k*-means (k = 8) clustering on the dynamically active regions (based on union differential H3K27Ac occupancy; |log_2_FC| ≥ 0.5, p_adj_ < 0.05 for all pairwise comparison). Cluster (C) 1 showed strong H3K27Ac and H3K4me1 signals on day 0 that are stably diminished as cells differentiate into memory cells and are coupled with the loss of ETS motifs. C2 included regions that are involved in NK cell innate effector function in response to pro-inflammatory cytokines, as exemplified by *Ifng* and *Cish*, among others. Coincidentally, this cluster is highly enriched in motifs for STAT proteins, which are critical for proinflammatory cytokine signaling in NK cells during early MCMV infection^11–15^. Indeed, pathways enriched in this cluster include cytokine, JAK/STAT, and T cell receptor (TCR) signaling pathways, which coincided with early innate effector functions on day 2 PI. Remarkably, the depositions of H3K27Ac and H3K4me1 in this cluster are highly coordinated and transient. Meanwhile, C3, C4 and C5 represent regions that are widely parallel and highly active during days 4, 7 and 14 PI, respectively, and are enriched for basic leucine zipper (bZIP) and RUNX motifs. Genes that are associated with these regions, such as *Nfkb1* and *Runx3*, as well as the transcription factor motifs found in these clusters have been implicated in NK cell maturation and effector responses (Fig. 1F, Supp. Fig. 2F)^16,17^. Furthermore, C6 are regions that transiently lost both histone marks from naïve cells but regained them in memory cells, highlighting shared regions between naïve and memory NK cells, whereas C7 and C8 delineate active and permissive regions that are enriched in memory cells and are associated with the adaptive immune system (Fig. 1F). Interestingly, although the trajectories of H3K27Ac and H3K4me1 in C8 are highly similar, there is a divergence between the two histone marks in C7 after day 14 PI, as H3K4me1 signals in these regions return to naïve baseline while H3K27Ac is still maintained at high levels (Supp. Fig. 2F). Transcription factor motifs found in C7 and C8 belong to the KLF and AP-1 family members, both of which have been implicated in the generation of immune memory^3,4,18^. Together, these data suggest that the multifaceted histone remodeling of NK cells in response to viral infection occurs in a coordinated (Spearman’s *ρ* = 0.50, *P* < 2.2 × 10^−16^, see Methods) stepwise manner and there are distinct waves of histone modifications that control innate and adaptive features of NK cells (Fig. 1E-F).

### Higher-order 3D chromatin topology in innate and adaptive lymphocytes is stable during infection

Epigenetic regulation of immune cells has been traditionally studied in the context of the 1D linear genome. However, the eukaryotic genome exists in a 3D space and is organized into multilayer hierarchical structures, namely chromatin compartments (hereafter referred to as A/B compartments that correspond to ‘active’ and ‘inactive’ compartments, respectively) and TADs^8,19^. However, less is known on how the 3D genome is changing as immune cells undergo extensive molecular adaptations in response to pathogens. Therefore, to determine how the chromatin of innate and adaptive lymphocytes are rewired beyond the 1D genome, we subjected both Ly49H^+^ NK cells and MCMV-specific CD8^+^ T cells at different time points following MCMV infection to HiC (Supp. Fig. 1, Supp. Fig. 3A-B)^19^. Although A/B compartments have been shown to undergo dynamic 3D reorganization during the development of various cell types^7,20^, it is unclear whether the A/B compartments of mature lymphocytes undergo similar changes during cell state transitions. To this end, we analyzed the A/B compartments from Ly49H^+^ NK cells sorted from day 0, 2, 7, and 35 PI, as well as naïve CD8^+^ T cells (day 0) and MCMV-specific M45^H2-Db+^ CD8^+^ T cells from day 7 and 35 PI. Additionally, we incorporated similar analysis on a previously published dataset generated from red pulp macrophages (RP macrophages) and plasmacytoid dendritic cells (pDCs) to universally understand how the higher-order chromatin compartments are organized between distinct immune cell types and cell states^21^.

**Figure 3.**
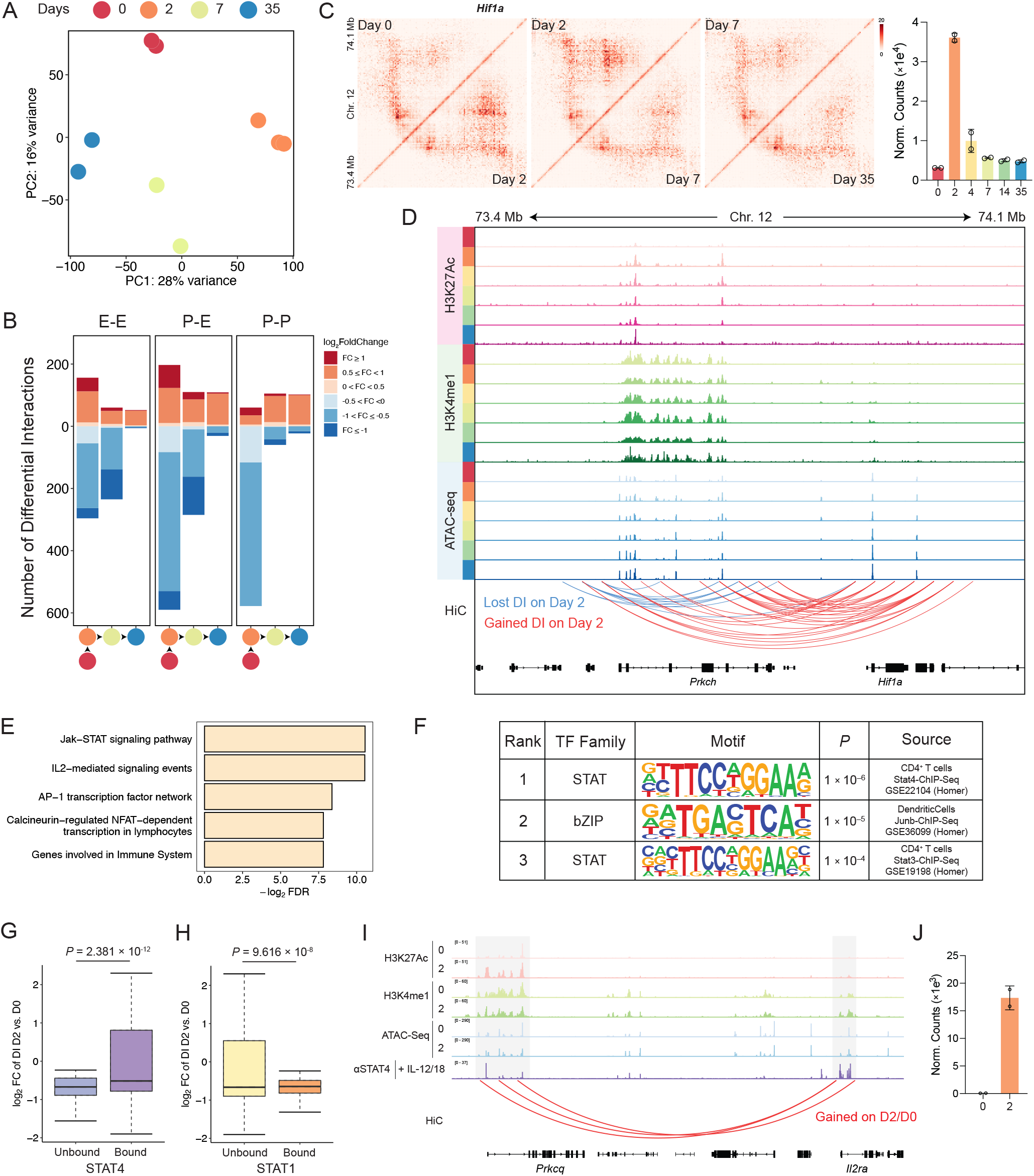
Innate lymphocytes undergo dynamic 3D chromatin remodeling during viral infection. (A) PCA of chromatin interactions in Ly49H^+^ NK cells during MCMV infection at 25kb bin resolution on day 0, 2, 7 and 35 PI. (B) Number of differentially interacting (DI) regions (p_adj_ <0.05) that either gain (red) or lose (blue) interactions at indicated transition time points categorized based on promoter-promoter (P-P), promoter-enhancer (P-E), or enhancer-enhancer (E-E) interactions. (C) Contact matrices heatmap at 5kb resolution of the *Hif1a* locus and its neighboring loci as Ly49H^+^ NK cells transition from d0 to d35 PI depicted as z-value (left). Gene label is located relative to the gene position in the genome. Normalized RNA counts of *Hif1a* transcript in Ly49H^+^ NK cells throughout infection. (D) Genomic tracks depicting H3K27Ac, H3K4me1, and ATAC-seq signals throughout different time points, as well as differentially interaction regions on d2 PI vs. d0 assessed by HiC that corresponds to the *Hif1a* locus in (C). (E) Top 5 most upregulated pathways on gained DI regions on d2 PI vs. d0 in Ly49H^+^ NK cells. (F) Known transcription factor motifs that are gained in DI regions on d2 PI vs. d0 in Ly49H^+^ NK cells. (G) Boxplot depicting the log_2_ fold change of DI regions in loops that are either bound or not bound by STAT4. *P*-value was calculated using two-sided student’s t-test. (H) Boxplot depicting the log2 fold change of DI regions in loops that are either bound or not bound by STAT1. *P-*value was calculated using two-sided student’s t-test. (I) Genomic tracks depicting H3K27Ac, H3K4me1, and ATAC-seq signals (d0 and d2 PI), and STAT4 ChIP-seq from *in vitro* stimulated NK cells, as well as gained DI regions on d2 PI vs. d0 in *Il2ra* locus. (J) Normalized RNA counts of *Il2ra* transcripts on d0 and d2 PI.

PCA of A/B compartment scores on Ly49H^+^ NK, CD8^+^ T, RP Macrophages and pDCs demonstrated that the compartmentalization of NK and CD8^+^ T cells was more similar to one another, and distant from the compartments of RP macrophages and pDCs (Fig. 2A), although cell type-specific differences are captured when these lymphocytes are isolated together (Supp. Fig. 4A). Strikingly, 93.8% of A/B compartments (48.5% of A compartment, and 45.3% of B compartment of all compartments) were shared between Ly49H^+^ NK cells and CD8^+^ T cells and were stable throughout the course of infection, and only 6.2% of their compartments were considered ‘dynamic’, which we defined as either distinct between the two cell types, or ‘flipped’ at least once during differentiation in either cell types (Fig. 2B). Of those that were ‘dynamic’, only about 20% were stable cell type-specific compartments, and about 34% compartments experienced cell type-specific switching during infection. Notably, over 10% flipped from B to A in CD8^+^ T cells during differentiation (Supp. Fig. 4B-C). Indeed, the average gene expression that spanned the ‘stable A’ compartments was higher compared to those that constituted ‘stable B’ regions (Supp. Fig. 4C). For example, the *Stat5a* and *Stat5b* loci, which are highly expressed and accessible in both cell types, fell under ‘stable A’ compartment. In contrast, cell type-specific genes, such as *Klf12* and *Themis*, showed a stable cell type-specific compartmentalization, consistent with the chromatin loop patterns within these loci (Fig. 2C, E). In contrast, the *Tbx21* locus, which encodes the transcription factor T-BET, was found to be part of ‘stable A’ compartment despite lacking expression and permissive epigenetic marks in naïve CD8^+^ T cells (Fig. 2D). A similar observation was found in the *Ifng* locus, an important effector cytokine that is expressed by NK cells, as well as effector and memory CD8^+^ T cells, but not by naïve cells (Supp. Fig. 4E). Thus, these data suggest that the chromatin compartmentalization of innate and adaptive lymphocytes are largely stable and may be established during development to poise for effector functions.

**Figure 4.**
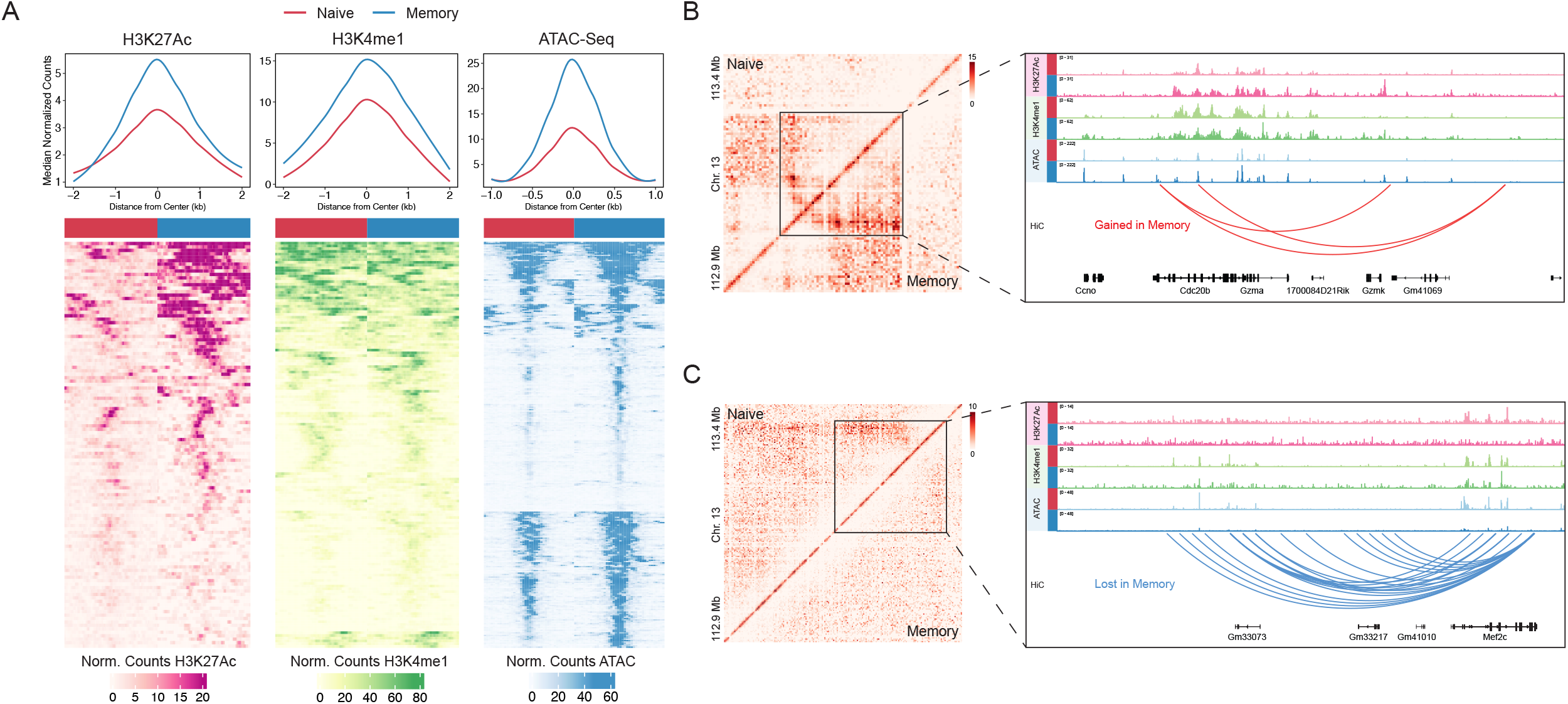
Memory NK cells are epigenetically poised at the 3D chromatin level. (A) Global H3K27Ac, H3K4me1, and ATAC-seq profiles of gained DI regions in naïve vs. memory Ly49H^+^ NK cells. (B&C) Contact heatmaps of the *Gzmk* (A), *Mef2c* (B) loci and their neighboring loci between naïve vs. memory Ly49H^+^ NK cells depicted as z-value at 5kb resolution and the accompanying genomic tracks featuring H3K27Ac, H3K4me1, ATAC-seq, and HiC DI regions.

We next analyzed TADs, which are fundamental units of the 3D genome that sequester ‘local’ regulatory element interactions to prevent aberrant gene expression^22^. We first identified TADs per sample and restricted our subsequent analyses on TADs that were reproducible between sample replicates (see Methods). To understand how TADs are changing during infection, we analyzed TADs within each cell type after infection and found that more than 68% and 60% of TADs were shared between at least two time points in Ly49H^+^ NK and CD8^+^ T cells, respectively, with the majority of TADs shared across all time points (Supp. Fig. 4D-E). To understand how TAD dynamics behave between cell types, we generated a cell type-specific TAD atlas by taking the union of TADs across all time points after infection from either Ly49H^+^ NK or CD8^+^ T cells. Consistent with our previous finding in A/B compartments, we also observed a high degree of TAD conservation between the two lymphocytes across different time points, with at least 52% of TADs shared across cell types throughout the course of infection, as demonstrated by TADs that demarcate the *Ifng* and *Ifng-as1* loci (Fig. 2F-G). Altogether, our data indicate that the majority of higher-order hierarchical structures of innate and adaptive lymphocytes are organized similarly and remain largely stable during infection.

### Chromatin interactions of naïve innate versus adaptive lymphocytes are distinct

Although NK cells and CD8^+^ T cells share a large proportion of higher-order 3D structures that are stable during infection, chromatin interactions (or loops) may inform an additional layer of regulation that is not captured by A/B compartment and TAD analyses. To identify chromatin loops, we used the HiC-DC+ algorithm to identify interactions within each sample at 25kb resolution (FDR <0.1), and created an atlas based on the union of interactions within each sample, which resulted in an atlas of 212,149 interactions at 25kb bin (Supp Fig. 3C)^23^. At the same time, we also quantified interactions at 5kb resolution to create a high-resolution map for contact matrix heatmap, but this resolution was not used for differential interaction (DI) analysis due to counts sparsity. From this 25kb bin interaction atlas, we performed DI analysis between naïve Ly49H^+^ NK and naïve CD8^+^ T cells. As discussed above, *Stat5a/b* showed a shared interaction pattern in NK and CD8^+^ T cells, whereas *Themis* and *Klf12* loci showed cell lineage-specific loops (Fig. 2E). Additionally, 26,410 DI regions (p_adj_ <0.05 & |log_2_FC| ≥0.5) were found to be distinct in naïve CD8^+^ T and NK cells (Supp. Fig. 5A). Motif analyses of accessible elements within DI regions between these two lymphocytes revealed enrichment of transcription factor motifs that belong to the HMG family members, such as *Tcf7* and *Lef1*, in CD8^+^ T cells, and T-BET (*Tbx21*) as well as AP-1 factors in NK cells (Supp. Fig. 5B). Indeed, these transcriptions factors have been implicated in regulating CD8^+^ T and NK cell development and functions^24,25^, and TCF7 itself has been shown to enforce chromatin interactions during T cell development by colocalizing with CTCF and recruiting NIPBL^20,26^.

**Figure 5.**
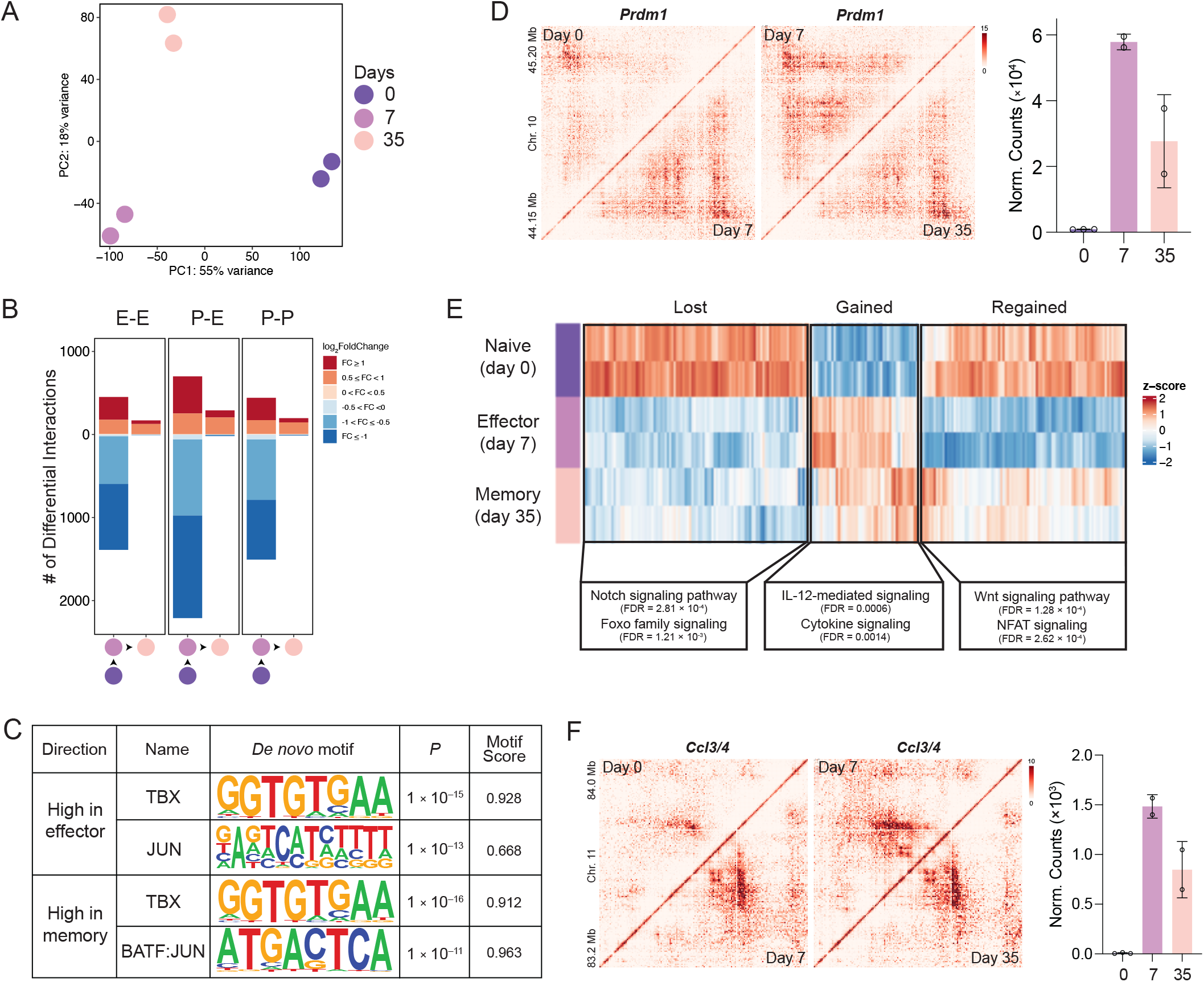
Memory CD8^+^ T cells maintain an effector-like 3D architectural program. (A) PCA of chromatin interactions in M45^H2-Db+^ CD8^+^ T cells during MCMV infection at 25kb bin. (B) Number of differentially interacting (DI) regions (p_adj_ < 0.05) that either ‘gain’ (red) or ‘lose’ (blue) interactions at indicated transition time points categorized based on promoter-promoter (P-P), promoter-enhancer (P-E), or enhancer-enhancer (E-E) interactions. (C) *De novo* transcription factor motifs that are enriched in effector and memory CD8^+^ T cells vs. naïve CD8+ T cells. (D) Contact matrices heatmap at 5kb resolution of the *Prdm1* locus and its neighboring loci as CD8^+^ T cells transition from d0 to d35 PI depicted as z-value (left). Normalized RNA counts of *Prdm1* transcript in CD8+ T cells throughout infection. Gene label is located relative to the gene position in the genome. (E) Heatmap of DI patterns in CD8^+^ T cells (*k*-means = 3) along with enrich pathways found within each cluster. (F) Contact matrices heatmap at 5kb resolution of the *Ccl3/4* locus and its neighboring loci as CD8+ T cells transition from d0 to d35 PI depicted as z-value (left). Normalized RNA counts of *Ccl3* transcript in CD8^+^ T cells throughout infection. Gene label is located relative to the gene position in the genome.

To validate this analysis, we sought to determine whether binding of TCF7 or T-BET can predict cell type-specific interaction in naïve cytotoxic lymphocytes. By overlapping DI regions between naïve NK and CD8^+^ T cells with either TCF7 ChIP-seq from naïve CD8^+^ T cells or T-BET ChIP-seq from naïve NK cells, we found that regions bound by TCF7 were biased towards CD8^+^ T cell-specific loops, while T-BET binding was predictive of NK cell identity (*P* <2.2×10^−16^ for all pairwise comparison) (Supp. Fig. 5C-D). Subsequently, of the DI regions, the most enriched pathways highlighted the differences in cell identities and relevant pathways for each cell type. For example, NK cell-mediated cytotoxicity and cytokine ligand-receptor interaction pathways were highly upregulated in NK cells, while various pathways involving TCR, WNT and Notch signaling were found to be higher in CD8^+^ T cells (Supp. Fig. 5E). Hence, the chromatin interactions between these naïve cell types are shaped by distinct families of transcription factors that mediate shared and/or cell type-specific 3D chromatin landscape.

### Innate lymphocytes undergo dynamic 3D chromatin remodeling during viral infection

Although NK cells undergo significant epigenetic and transcriptional remodeling during naïve to memory transition, it remains unclear whether these changes are accompanied by active reorganization of 3D chromatin loops. PCA of the interaction atlas in differentiating NK cells revealed distinct chromatin interaction profiles at different time points after infection (Fig. 3A). The transition from day 0 to 2 PI showed the greatest transitional changes, with 3994 DI regions showing significant changes (p_adj_ <0.05) of varying magnitudes (Fig. 3B and Supp. Fig. 6A). Moreover, promoter-enhancer (P-E) as well as enhancer-enhancer (E-E) interactions showed the strongest changes (|log2FC| ≥1) compared to promoter-promoter (P-P) interactions (Fig. 3B). *Hif1a* was among the most significant DI regions, and we observed gain of proximal interactions of the *Hif1a* locus with its neighboring gene, *Prkch*, and simultaneous loss of *Prkch* local self-interactions that correlated with an increased in *Hif1a* expression, but not with *Prkch* expression, on day 2 PI (Fig. 3C-D and Supp. Fig. 6B). Additionally, these correlates coincided with increased H3K27ac occupancy and chromatin accessibility in an already highly permissive *Prkch* locus, and losing these proximal loops on day 7 PI was associated with a drop in *Hif1a* transcripts (Fig 3D). Furthermore, we observed other interaction features in which the loss or gain of distal loops may or may not be correlated with gene expression of the interacting loci. For instance, the loss of distal interactions between *Gzmb* and *Cma1* is accompanied by a decrease and a concurrent increase in H3K27Ac occupancy in *Cma1* and *Gzmb* loci, respectively, and is reflective of the transcript levels of both genes (Supp. Fig. 6C). In contrast, gain of distal interactions between the *Ifng* regulatory elements with *Mdm1* on day 2 PI only correlates with *Ifng* expression (Supp. Fig. 6D). Altogether, our data highlight how proximal and distal epigenomic changes may influence the transcription of surrounding regions through a 3D mechanism.

**Figure 6.**
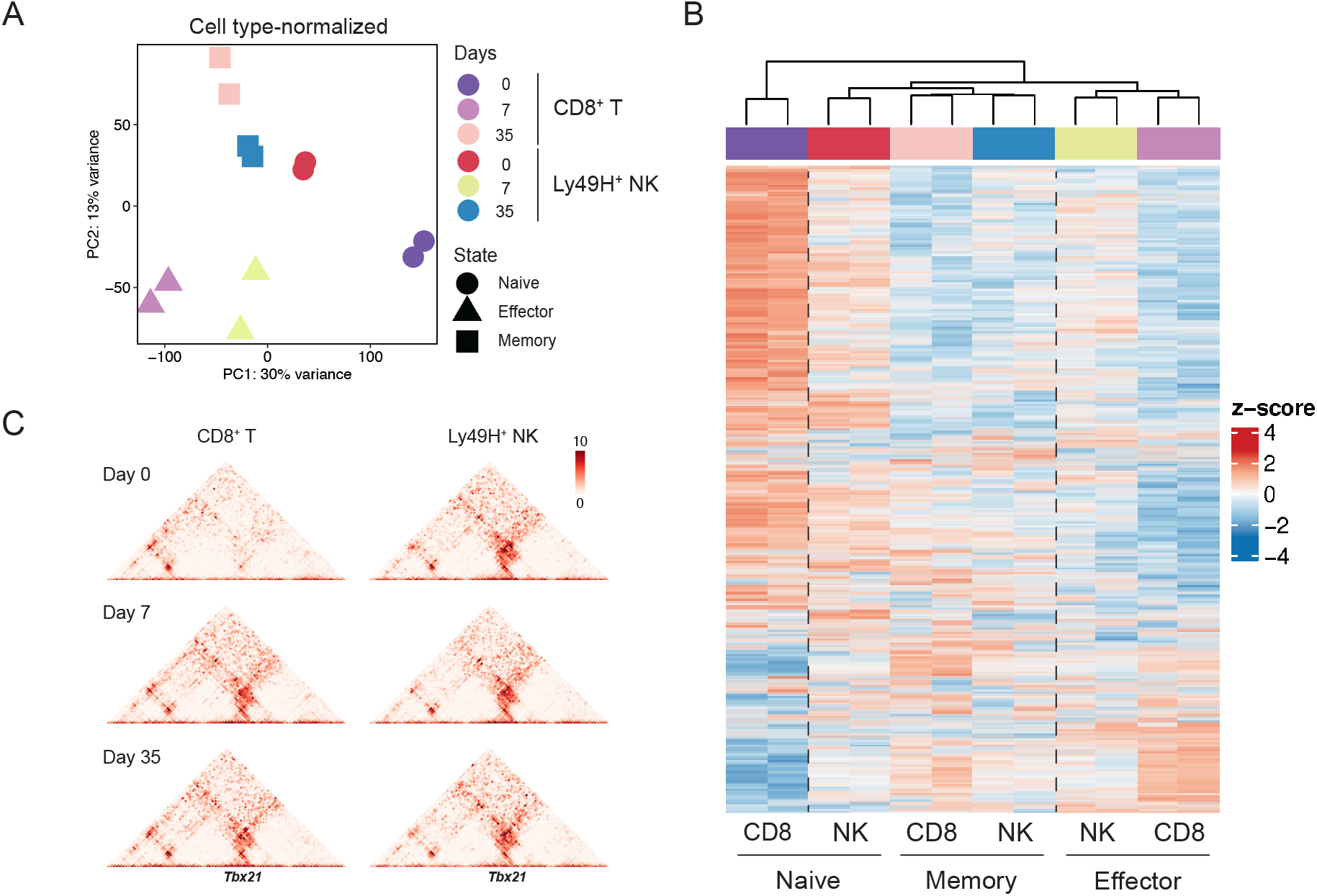
Effector and memory CD8+ T cells adopt NK-like three-dimensional chromatin architecture. (A) Cell type-normalized PCA of Ly49H^+^ NK cells and M45^H2-Db+^ CD8^+^ T cells following MCMV infection at 25kb bin. (B) Heatmap of mean-centered log_2_ normalized counts of chromatin loops with the greatest variance. (C) Contact heatmap matrices at 5kb resolution on the *Tbx21* locus on day 0, 7, and 35 PI in both CD8^+^ T cells and Ly49H^+^ NK cells. Gene label is located relative to the gene position in the genome.

Given that the 3D chromatin loops of Ly49H^+^ NK cells are most dynamic early after infection, we performed pathway enrichment analysis of highly DI regions that were gained on day 2 PI compared to day 0. Our analysis revealed that the top 5 most enriched pathways were dominated by NK cell responses to cytokine and antigen signaling (Fig. 3E), along with the transcription factors that may be regulating these processes, such as JAK/STAT, AP-1 and NFAT (Fig. 3E-F). To test whether STAT binding is associated with differential interactions, we overlapped DI regions from the day 2 PI versus day 0 comparison with STAT4- or STAT1-bound peaks from *in vitro* IL-12+IL-18 or IFN-*α* stimulated NK cells, respectively^17,27^. We found that loops bound by STAT4 showed increased interactions on day 2 PI compared to those that were not bound by STAT4 (*P* = 2.381 × 10^−12^) (Fig. 3G). This type of interaction is exemplified by the *Il2ra* locus, which exhibited a similar interaction pattern and epigenomic remodeling to the *Hif1a* locus (Fig. 3C-D, I-J). In contrast, loops that were bound by STAT1 showed a slight but significant decrease in interactions (*P* = 9.616 × 10^−8^) as illustrated by the *Prkcq* and *Irf2* loci (Fig. 3H, Supp. Fig. 6E-G). Thus, Ly49H^+^ NK cells experience a substantial rewiring of the 3D chromatin interactions throughout the course of infection, and STAT family members may be involved in coordinating early looping events that shape the overall NK cell response to viral infection.

### Memory innate lymphocytes are epigenetically poised at the 3D chromatin level

We have previously demonstrated that memory NK cells are epigenetically distinct from naïve NK cells and may be poised to be more responsive to secondary challenge via differences in chromatin accessibility^3^. In addition, PCA of differentiating NK cells suggests that the 3D chromatin landscape of memory Ly49H^+^ NK cells is unique from naïve NK cells, with 858 loops found to be DI (Fig. 3A and Supp. Fig. 6A). To investigate the relationship between the 1D epigenetic changes and the 3D chromatin reprogramming, we integrated histone profiling and chromatin accessibility on DI loops in memory versus naïve Ly49H^+^ NK cells. Globally, loops that were gained in memory cells also showed higher levels of H3K27Ac and chromatin accessibility, and to a lesser extent H3K4me1 (Fig. 4A). For example, among those that were gained in memory cells, the neighboring *Gzma, Gzmk* and *Cdc20b* loci showed higher expression in memory cells and were epigenetically more permissive compared to naïve Ly49H^+^ NK cells (Fig. 4B and Supp. Fig. 6H). In contrast, *Mef2c*, a transcription factor downstream of MAP kinase signaling pathway, showed significant downregulation of permissive histone marks and chromatin accessibility, which corresponded to the loss of interactions and transcription in this locus (Fig. 4C and Supp. Fig. 6I). Therefore, epigenetic poising of the memory state in innate lymphocytes not only occurs in 1D, but also in 3D.

### Memory CD8^+^ T cells maintain an effector-like 3D architectural program

Whereas naïve NK cells can readily exert their effector functions in response to various stimuli, naïve CD8^+^ T cells are less responsive and more ‘quiescent’ compared to their innate counterparts^28^. Moreover, the differences in their reactivity can partly be explained by the differences in the epigenetic states of these cells. Indeed, previous studies have shown that many effector genes in naïve CD8^+^ T cells are kept in bivalent states, while the chromatin of naïve NK cells are more ‘poised’ for effector functions^29,30^. Given that these innate and adaptive lymphocytes share many features and undergo parallel epigenetic and transcriptomic programming, we sought to determine how the 3D chromatin loops of differentiating CD8^+^ T cells are changing during viral infection. To this end, PCA was performed on HiC of naïve (day 0), effector (day 7 PI), and memory (day 35 PI) MCMV-specific M45^H2-Db+^ CD8^+^ T cells (Fig. 5A). PC1 revealed that naïve-to-effector T cell differentiation experienced the greatest changes in chromatin loops (Fig. 5A), with 18,149 loops changing in the naïve versus effector states, and only 1,386 loops distinct between effector and memory cells (Supp Fig. 7A). Interestingly, more of the DI loops were lost as cells differentiate from naïve to effector rather than gained, and the numbers of high-fold change (|log2FC| *≥*1) loops were substantial across the different types of interactions, with P-E interactions showing the greatest number of changes (Fig. 5B). Strikingly, the 3D chromatin of day 35 PI memory CD8^+^ T cells was more similar to day 7 PI effector cells than to naïve cells, and memory cells almost exclusively gained in DI compared to effector cells (Fig. 5A-B).

*De novo* transcription factor motif analysis of DI regions between naïve versus effector cells, and naïve versus memory cells, also suggested an enrichment of T-BOX family members (such as T-BET and/or EOMES) and AP-1 factors (namely JUN and BATF) (Fig. 5C). Consistently, among gene loci that gained DIs in effector cells and were stably maintained in memory cells, many are involved in the effector differentiation and function of CD8^+^ T cells. For example, *Prdm1*, a T-BET-regulated gene that encodes for the key transcription factor BLIMP-1 for driving effector CD8^+^ T cell differentiation, exhibited a dramatic gain of DIs that remained in memory cells despite the decreased in *Prdm1* transcript in memory cells compared to effector cells (Fig. 5D). Moreover, this stably ‘gained’ group was highly enriched in lymphocyte responsiveness to cytokines, as highlighted by the effector gene loci *Ccl3/Ccl4* and *Gzma/Gzmk* (Fig. 5E-F, Supp Fig. 7B). Hierarchical clustering analysis also identified two additional patterns of interaction dynamics in differentiating lymphocytes; those that were stably lost in effector and memory cells compared to naïve cells, and transient interactions that were lost from naïve to effector cells but regained in memory cells (Fig. 5E). In the stably ‘lost’ regions, we found that the Notch signaling pathway was particularly enriched, whereas regions involved in Wnt signaling seemed to be ‘regained’ in memory cells, as illustrated by the *Mapk9* and *Lef1* loci, respectively (Supp. Fig. 7C-D). Hence, these data suggest that memory CD8^+^ T cells adopt an effector-like 3D architectural program that may be driven by T-BET and AP-1 factors.

### Effector and memory CD8^+^ T cells adopt an NK-like 3D chromatin architecture

Lastly, to understand the 3D chromatin dynamics of Ly49H^+^ NK cells and CD8^+^ T cells as they differentiate from naïve to memory in parallel, we generated a combined interaction atlas together to directly compare the two cell types. Initial PCA indicated cell type-specific separation (Supp. Fig. 3D); however, from this initial analysis, it was apparent that effector and memory CD8^+^ T cells clustered closer to NK cells, compared to naïve CD8^+^ T cells. To account for cell type-specific differences, we used cell type-normalized values (see Methods) that may capture cellular states trajectory in response to infection more accurately. By doing so, we could observe that naïve NK cells clustered closely to memory CD8^+^ T cells (relative to other CD8^+^ T cell subsets), and memory NK cells were found in between memory CD8^+^ T cells and naïve NK. Meanwhile, day 7 PI effector NK cells and CD8^+^ T cells were found together and separated from the other clusters (Fig. 6A). Indeed, hierarchical clustering analysis from cell-normalized values confirmed that the 3D chromatin interactions of naïve CD8^+^ T cells were most distinct, as naïve NK cells are clustered together with memory NK and CD8^+^ T cells (Fig 6B). This clustering is exemplified by the *Tbx21* locus, where the interactions in effector and memory CD8^+^ T cells are comparable to NK cells at all time points, despite loss of T-BET expression in memory CD8^+^ T cells (Fig. 6B-C, Supp. Fig. 7E). Altogether, our data indicate that both innate and adaptive lymphocytes undergo parallel 3D genome remodeling as cells differentiate from naïve to effector to memory states, with CD8^+^ T cells experiencing greater overall changes as they adopt an NK-like architectural fate that may explain their rapid responsiveness during reactivation.

## DISCUSSION

Accumulating evidence in the past decade have shown that NK and CD8^+^ T cells possess many similar features in response to viral infection that can be partly attributed to parallel transcriptomic and 1D epigenetic remodeling between the two related cell types^3,5^. In this study, we define the 3D chromatin landscape dynamics during innate and adaptive immune memory acquisition. We establish that innate and adaptive lymphocytes undergo substantial 3D genomic reprogramming as they encounter viral infection, with a decoupling of higher-order structure from finer-scale chromatin looping. In addition, our results also suggest the importance of a 3D mechanism beyond the 1D epigenetic state in controlling gene expression in differentiating lymphocytes that experience extensive molecular adaptations. We also highlight distinct families of transcription factor networks that may be required during stage-specific transitions. And lastly, we observe that these antiviral innate and adaptive lymphocytes undergo parallel 3D epigenetic imprinting during their differentiation from naïve to memory cells.

Our understanding of the role of the 3D genome in regulating immunity is only in its infancy, with most studies focusing on the dynamics of 3D chromatin in the context of immune cell development or acute innate inflammatory responses. Within the larger hierarchical structures, we found that innate and adaptive lymphocytes share high proportions of compartments and TADs that are largely stable throughout effector and memory differentiation. In support of our findings, previous studies in lymphocyte development suggest that higher-order structures are established as early as the common lymphoid progenitor (CLP) stage, with relatively little changes occurring after the CLP and during acute inflammatory response, consistent with the idea that the higher-order structures may be acting as a scaffold to facilitate cell type-specific functions^7,20,31,32^. However, how these larger structures are established during lymphocyte development and antiviral responses, and what factors shape the higher-order structures within innate and adaptive lymphocytes, warrant further investigation.

Although the higher-order structures of antiviral NK cells and CD8^+^ T cells are largely stable, our chromatin interaction analyses suggest dynamic locus-specific reprogramming that are likely to be mediated by distinct transcription factor families during infection. In innate lymphocytes, the cytokine signaling networks are mediated by stimulus-dependent STAT family members that are induced early during infection. Interestingly, our data suggest opposite roles of STAT1 and STAT4 in mediating chromatin interactions, consistent with their previously described antagonism of one another^15^. Compared to NK cells that are enriched in AP-1 and TBX motifs, chromatin interactions in naïve CD8^+^ T cells appear to be partly mediated by HMG transcription factor family members such as TCF7 and LEF1, which have been shown to enforce CD8^+^ T cell chromatin architecture and identity. However, as differentiating CD8^+^ T cells adopt an NK-like chromatin architecture in effector and memory cells, they may similarly require AP-1 and TBX factors. Indeed, BATF, a member of the AP-1 family, has been shown to recruit the structural protein CTCF in effector CD4^+^ T cells^33^; however, it remains unclear how the T-box transcription factors are modulating these processes. Understanding the interplay between T-box, TCF, and AP-1 factors in regulating the 3D chromatin architecture during effector and memory differentiation will offer valuable insights into the intricacies of the higher order 3D regulation of NK and CD8^+^ T cells responding to viral infection.

In summary, we provide evidence for 3D chromatin reprogramming in innate and adaptive lymphocytes during immunological memory acquisition, and generate a 3D genome atlas of these cells responding to a viral infection. These findings will collectively expand our understanding of how regulatory element networks orchestrate gene expression, and will provide insights into how non-coding genetic elements control optimal immune responses. In addition, our atlas may inform how and why suboptimal or hyperinflammatory immune responses occur, as in the case of primary immunodeficiencies, autoimmunity, and inflammatory disease. It has become increasingly evident that many non-coding genetic polymorphisms are associated with various diseases^34–37^, and future studies will connect the specificity and the interactions of these regulatory elements with disease initiation and progression^38,39^. Thus, integration of tiered genetic information, including the epigenetic state, transcriptome, and 3D spatial chromatin landscape, can be harnessed for both diagnostic purposes and the development of therapy for various genetic disorders and diseases.

## MATERIALS & METHODS

### Animals

All mice used in this study were housed and bred under specific pathogen-free conditions at Memorial Sloan Kettering Cancer Center and handled in accordance with the guidelines of the Institutional Animal Care and Use Committee (IACUC). The following mouse strains were used in this study: C57BL/6 (CD45.2), C57BL/6-CD45.1 (CD45.1), *Klra8*^*−/−*^ CD45.1xCD45.2 (Ly49H-deficient). Experiments were conducted using 8-to 12-weeks old mice and experiments were done using age- and gender-matched mice in accordance with approved institutional protocols.

### Adoptive transfer and viral infection

MCMV (Smith strain) was serially passaged through BALB/c hosts three times, and then salivary gland viral stocks were prepared with a homogenizer for dissociating the salivary glands of infected mice 3 weeks after infection. For adoptive transfer studies, approximately 10^5^ NK cells from WT (CD45.1 or CD45.2) mice were injected into Ly49H-deficient recipients 1 day prior to MCMV infection. Recipient mice in adoptive transfer studies were infected with MCMV by i.p. injection of 1.1 × 10^3^ plaque-forming units (PFU) in 0.5 ml of PBS. For direct infection experiments, WT mice were infected with 1.1 × 10^4^ PFU in 0.5 ml of PBS.

### Isolation of lymphocytes, flow cytometry and cell sorting

Spleens were dissociated with glass slides and filtered through 100-μm cell strainer. Flow cytometry and cell sorting were done on the Cytek Aurora (Cytek Biosciences) and Aria II cytometers (BD Biosciences), respectively. Before cell sorting, NK cells were enriched by incubating whole splenocytes with the following antibodies at 20 μg/ml against CD3ε (17A2), CD4 (GK1.5), CD8 (2.43), Ter119 (TER-119), CD19 (1D3), Ly6G (1A8) (BioXCell) followed by magnetic depletion using goat anti-rat beads (QIAGEN). For adoptive transfer experiments, enriched splenocytes were then stained with surface markers to identify Ly49H^+^ NK cells (CD3/TCRb/CD19^−^ NK1.1^+^ CD49b^+^ Ly49H^+^) and sorted based on congenic markers (CD45.1 and CD45.2) to identify donor cells. Naïve CD8^+^ T cells (CD8a^+^ CD3/TCRb^+^ CD62L^+^ CD44^−^), and MCMV-specific CD8^+^ T cells (CD19/NK1.1^−^ CD8a^+^ CD3/TCRb^+^ CD44^+^ M45^H2Db+^) were collected from directly infected WT mice on day 0, 7, and 35 post infection. In both adoptive transfer and direct infection setting, cells were sorted with >95% purity. For intracellular staining, cells were surface stained, fixed with eBioscience™ Intracellular Fixation & Permeabilization Buffer for 30 mins at 4^°^C, and stained with T-bet antibody for 30 mins in 1X eBioscience™ Permeabilization Wash Buffer.

### High-throughput chromatin capture assay (HiC

Chromatin capture assay was done according to Arima HiC kit protocol. Briefly, 1.5-2 million cells per replicate were sorted, crosslinked, and digested with restriction enzyme cocktail. Digested DNA overhangs were then repaired and filled in with biotinylated nucleotides and re-ligated to produce proximal-ligated DNA that can be sheared to enrich for ‘chimeric’ pairs that contain biotinylated fragments. DNA libraries were then prepared using KAPA HyperPrep Kit with an average amplification of 5 cycles. After quality control by Agilent BioAnalyzer and PicoGreen quantification when applicable, libraries were pooled equimolar and run on NovaSeq6000 or HiSeq4000 in a PE100 run, using the NovaSeq6000 S2 or S4 Reagent Kit (200 cycles) or HiSeq3000/4000 SBS Kit (Illumina). The loading concentration was 0.5 nM (NovaSeq) or 3-3.8nM (HiSeq) and a 1-5% spike-in of PhiX was added to the run to increase diversity and for quality control purposes. Samples were first ‘shallow’ sequenced to assess the quality of the sample. Upon quality control assessment using parameter described by Rao *et al*., the libraries were further sequenced^40^. For each sample, the runs yield on average about 680 million reads.

### HiC data processing

Fastq files were processed using the default parameters of Juicer LSF pipeline (v1.5.6) (github.com/aidenlab/juicer) with slight modifications to adjust for MSKCC’s LSF system (github.com/endicytosis), and BWA software (v0.7.17) was used for alignment of chimeric read pairs to mm10 reference genome to generate inter30.hic files. These inter30.hic files were used for subsequent downstream analyses using HiC-DC+ pipeline (github.com/mervesa/HiCDCPlus)^23^.

### A/B chromatin compartment and TAD analysis

Compartment analysis was done on Knight-Ruiz normalized inter30.hic files using the eigenvalue function from Juicer on each ‘shallow’ sequenced sample at 100kb bin. The eigenvalue direction is arbitrary, and thus we defined A/B compartments based on correlation with cell type specific ATAC values where positive and negative values were considered as A and B compartments, respectively. For further analysis, we filtered out compartment regions that do not share the same direction between replicates and performed ANOVA on the eigenvalues from each bin. ‘Stable’ A and B compartments are defined as regions that do not change directions between cell types and/or during infection, and ‘dynamic’ regions are identified as any regions that flipped direction at least once during infection regardless of cell types. For domain level analyses, inter30.hic files were ICE normalized at 50kb bin, and domains were called using ‘gi_list_topdom’ command at 50kb resolution with a window size of 10 for each sample^41^. We defined ‘high confidence’ TADs if they differ only by 50kb (one bin) and are reproducible across sample replicates.

### Chromatin loops analysis

Chromatin interactions/loops were called using HiC-DC+ package on inter30.hic files at 25kb resolution with FDR <0.1 and chromatin interaction atlas was generated by taking the union of identified loops from each sample. Differential interaction (DI) analysis was performed using ‘hicdcdiff’ command from HiC-DC+ package with ‘local’ fitType, granularity of 25,000, D_min_ and D_max_ of 0 and 2e6, respectively. Regions are considered differentially significant when p_adj_ <0.05. For cell type-specific analysis, we created an atlas by isolating interactions that were only found in that specific cell type. For cell type-normalized PCA, we applied ‘removeBatchEffect’ command from limma (v3.50.1) to the normalized log_2_ values using cell type as batch parameter^42^. To generate contact matrix heatmap, we called interactions at 5kb resolution and converted the counts to negative binomial Z-score normalized counts (z-value) and stored them as .hic files using ‘hicdc2hic’ command. All contact matrix heatmaps were plotted using plotGardener package (v3.16)^43^.

### Histone CUT&RUN

50,000 sorted cells were washed with PBS and resuspended in Buffer 1 (1X eBioscience Perm/Wash Buffer, 1X Roche cOmplete EDTA-free Protease Inhibitor, 0.5 uM Spermidine in H_2_O), and were incubated with control IgG, H3K4me1 (Epicypher), H3K27Ac (Epicypher) antibodies at 1/100 dilution in Antibody Buffer (Buffer 1 + 2uM EDTA) in 96 well V-bottom plate at 4^°^C overnight. Upon antibody incubation, cells were washed twice with Buffer 1 and resuspended in 50ul of Buffer 1 + 1X pA/G-MNase (Cell Signaling) and incubated on ice for 1 hr and washed twice with Buffer 2 (0.05% w/v Saponin, 1X Roche cOmplete EDTA-free Protease Inhibitor, 0.5 uM Spermidine in 1X PBS) three times. After washing, Calcium Buffer (Buffer 2 + 2uM of CaCl_2_) was used to resuspend the cells for 30 mins on ice to activate the pA/G-MNase reaction, and equal volume of 2X STOP Buffer (Buffer 2 + 20uM EDTA + 4uM EGTA) was added along with 0.1pg/ml of *E. coli* spike-in DNA. Samples were incubated for 15 mins at 37^°^C and DNA was isolated and purified using Qiagen MinElute Kit according to manufacturer’s protocol and subjected to library amplification. Briefly, DNA was quantified by PicoGreen and the size was evaluated by Agilent BioAnalyzer. Illumina sequencing libraries were prepared using the KAPA HTP Library Preparation Kit (KAPA Biosystems #KK8234) according to the manufacturer’s instructors with <0.001-3.8 ng input DNA and 14 cycles of PCR. Barcoded libraries were run on the NovaSeq6000 in a PE100 run, using the NovaSeq6000 S4 Reagent Kit (200 cycles) (Illumina). An average of 9 million paired reads were generated per sample.

### CUT&RUN data processing and analysis

Paired reads were trimmed for adaptors and removal of low-quality reads by using Trimmomatic (v0.36) and aligned to the mm10 reference genome using Bowtie2 (v1.2.3). Upon alignment, peaks were called using MACS2 (v2.1.1.20160309) with IgG control as ‘input’ using narrow peak parameters -p 1e-3 -n not_masked –keep-dup all –SPMR –nomodel^44^. After peak calling, bottom 25% of peaks of each sample was filtered out from the analysis, and only peaks that were shared between at least two replicates and overlapped with ATAC peaks are retained for further analysis. Peaks within ± 1000 bp away from each other were merged and considered as one peak. To assess the relationships between H3K27Ac and H3K4me1, we generated an overlapping peak atlas between the two histone marks and used this atlas for mapping and counting with the summarizeOverlaps function from GenomicAlignment package(v1.29.0)^45^. Differential occupancy analysis was done using DESeq2 (v.1.34.0)^46^. Promoter regions were defined as peaks that overlapped a region that was +2.5kb to −0.5kb from the transcriptional start site (TSS), while enhancer regions were defined using H3K4me1 that were not annotated as promoters.

### RNA-seq, ChIP-seq and ATAC-seq

Fastq files quality was assessed using FastQC (v0.11.7) (Babraham Bioinfomatics). Paired reads were trimmed for adaptors and removal of low-quality reads using Trimmomatic (v0.36). RNA-seq, ATAC-seq and STAT ChIP-seq (GSE106139) were processed as previously described^3,15^. T-BET ChIP-seq on NK cells (GSE145299) and TCF7 ChIP-seq on naïve CD8^+^ T cells (GSE164670) were aligned to the mm10 reference genome with Bowtie2 (v1.2.3). MACS2 (v2.1.1.20160309) was used to call peaks using parameters mentioned above^44,47,48^. For TCF7 ChIP-seq peak calling, Tcf7^−/−^ CD8^+^ T cells were used as an input control which resulted in 3,360 peaks. For T-BET ChIP-seq, the two replicates were subjected to Irreproducibility Discovery Rate (IDR) (v2.0.4.2) upon peak calling and further filtered using IDR value <0.01 to generate 6,000 surviving T-BET peaks.

### Transcription Factor Motif Analysis

All motif analyses were done using Hypergeometric Optimization of Motif EnRichment algorithm (HOMER v4.10)^49^ using direction-agnostic chromatin accessible atlas previously generated^3^, but filtered on cell type-specific ATAC peaks for cell type-specific analysis. Briefly, all identified CUT&RUN or HiC DI regions were overlapped with chromatin accessibility atlas and overlapping chromatin accessible peaks were used as inputs for HOMER with parameter “-size given -len 6,8,10,12 -mset vertebrates -mask” for *de novo* motif analysis. For HiC, chromatin accessible regions must overlap with at least one of the interacting anchors. For all analysis, a combined NK and CD8^+^ T cell ATAC-seq atlas, or cell type-specific atlas, were used as background.

### Pathway analysis

For histone CUT&RUN peaks, bed files of peak regions were used as input, and for HiC, bed files containing a combination of first and second anchor regions were used as input. These bed files were then subjected to analysis with rGREAT(v1.26.0)^50^ using default settings and MSigDB Pathway databases. Enriched pathways were then filtered using qvalue <0.05 cutoff as calculated by GREAT.

### Correlation analysis

Reproducibility of CUT&RUN data was performed by Pearson’s correlation analyses between samples on log_2_ normalized values matrices from each sample. To assess the relationship between H3K27Ac and H3K4me1, Spearman’s correlation was done on mean-centered log_2_ normalized matrices of the two histone marks from the overlapping atlas.

### Data and code availability

All data generated and supporting the findings of this study either have been deposited at GEO or will be deposited at GEO upon publication of the manuscript. GEO accession numbers are available within the paper (see Materials & Methods). General codes for analyses will be made available through github.com/endicytosis.

## ACKNOWLEDGEMENTS

We thank members of the Sun lab, and Drs. Steven Josefowicz and Lewis Lanier for critical feedback on this study, and the NIH Tetramer Core for providing reagents. We acknowledge the use of Integrated Genomics Operation Core, funded by the NCI Cancer Center Support Grant (CCSG, P30 CA08748), Cycle for Survival, and the Marie-Josee and Henry R. Kravis Center for Molecular Oncology. We also acknowledge the use of High-Performance Computing Group at Memorial Sloan Kettering. J.C.S. was supported by the Ludwig Center for Cancer Immunotherapy, the American Cancer Society, the Burroughs Wellcome Fund, and the NIH (AI100874, AI130043, AI155558, and P30CA008748).

## Author Contributions

E.K.S. and J.C.S. designed the study. E.K.S performed experiments and bioinformatics analyses. C.M.L and M.S performed bioinformatics analyses. C.S.L and J.C.S provided critical resources and supervision for the study. E.K.S. and J.C.S. wrote the manuscript.

## SUPPLEMENTARY FIGURE LEGENDS

**Supplementary Figure 1.**
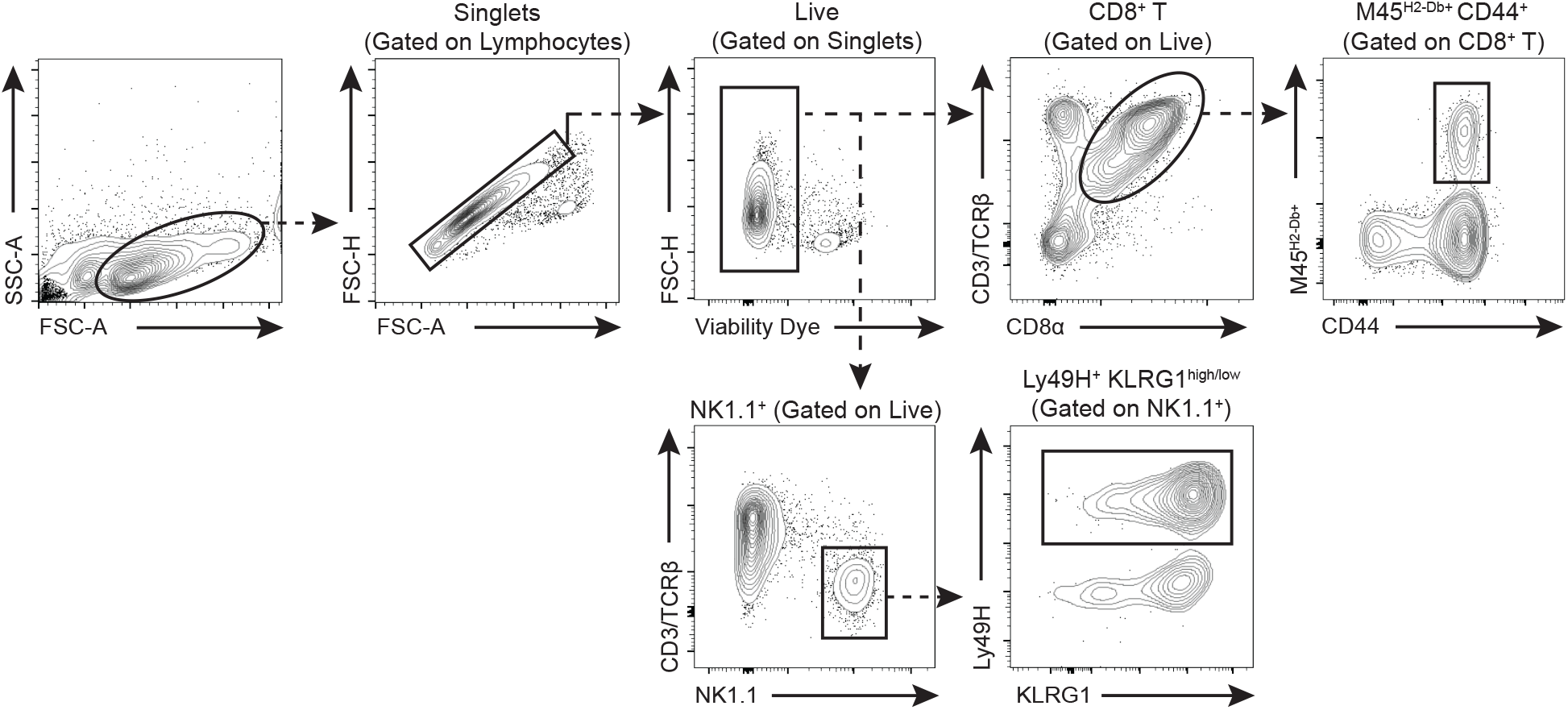
Cell sorting strategy of MCMV-specific Ly49H^+^ NK and M45^H2Db+^ CD8^+^ T cells.

**Supplementary Figure 2.**
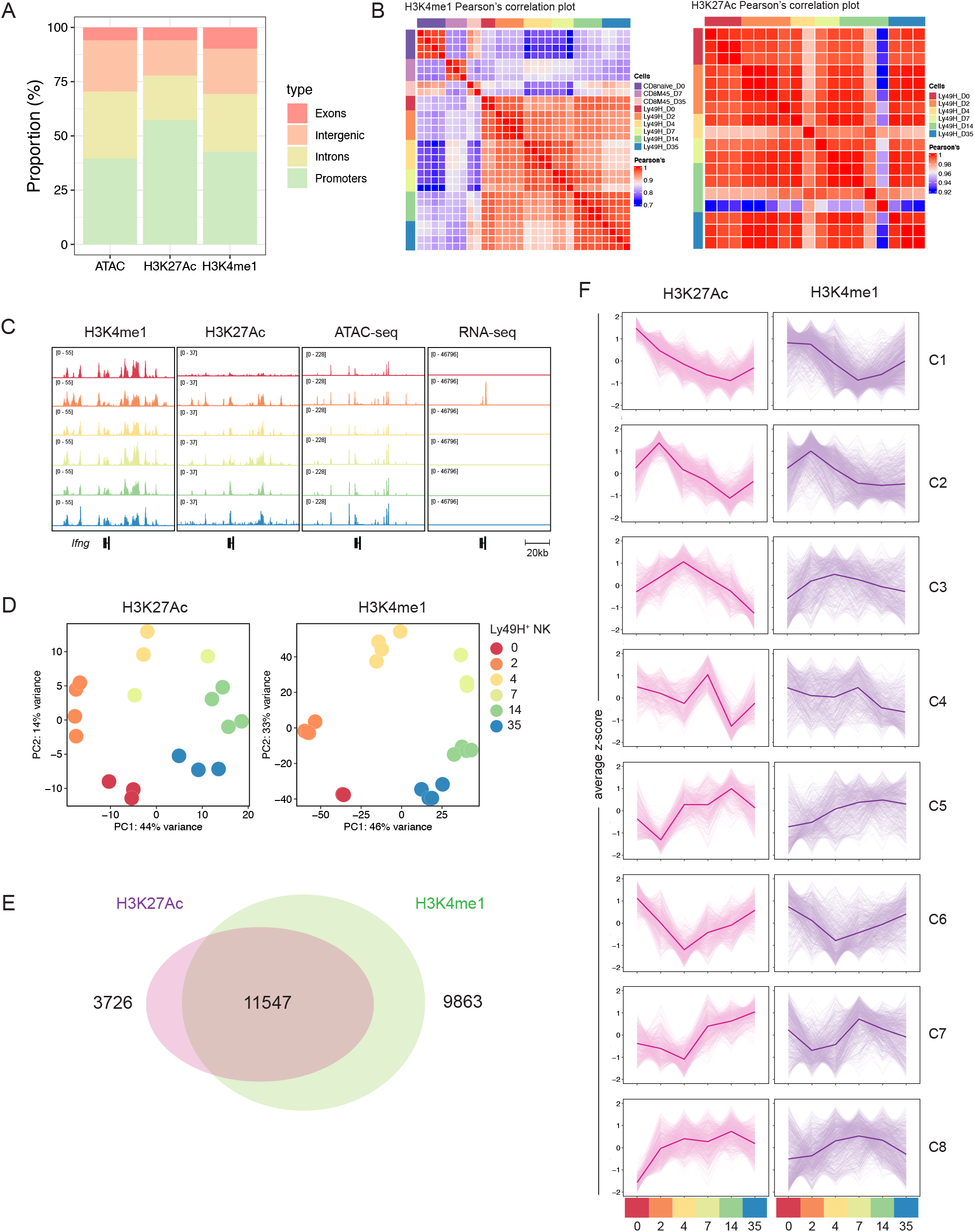
CUT&RUN analysis and quality control. (A) Annotated peak distributions between ATAC, H3K4me1, and H3K27Ac atlas in Ly49H^+^ NK cells during MCMV infection. (B) Correlation plot of H3K4me1 (top) and H3K27Ac (bottom) for each sample replicate analyzed separately. (C) H3K4me1, H3K27Ac, ATAC-seq, and RNA-seq tracks in the *Ifng locus* of Ly49H^+^ NK cells at different time points post MCMV infection. (D) PCA analysis of Top 5000 H3K4me1 regions in Ly49H^+^ NK cells (top), and top 500 H3K27Ac regions in Ly49H^+^ NK cells (bottom). (E) Venn diagram of H3K4me1 and H3K27Ac overlapping peaks that were used to assess the relationship between the two histone modifications (Figure 1E). (F) Mean z-score similar to Fig. 1E but depicted as line plots.

**Supplementary Figure 3.**
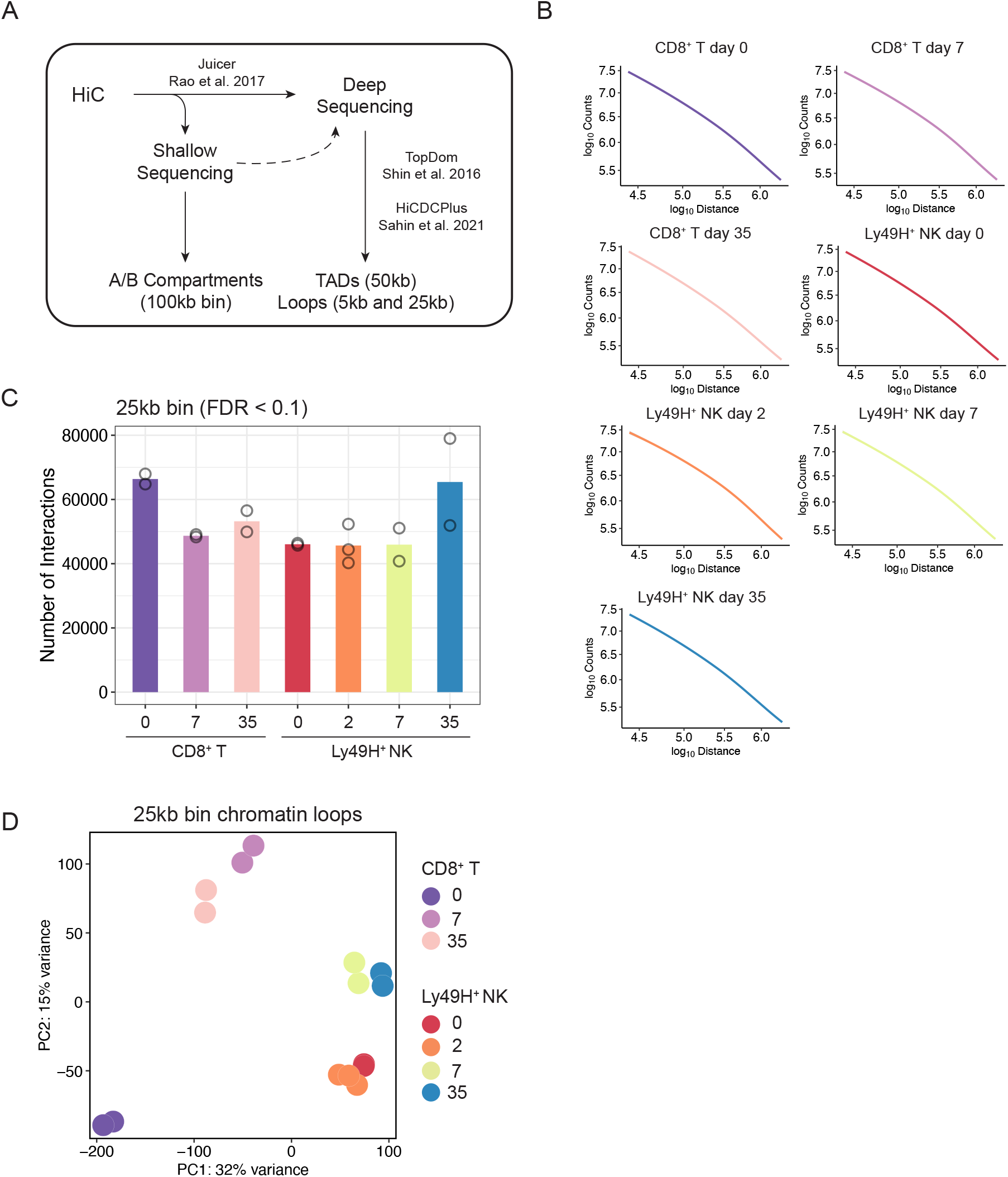
HiC analysis pipeline and quality assessment. (A) Schematic of HiC analysis pipeline. (B) Interaction counts as a function of genomic distance in each sample. (C) Number of significant interactions called by HiCDC+ in each sample. (D) PCA of union of significant interactions in all samples analyzed together.

**Supplementary Figure 4.**
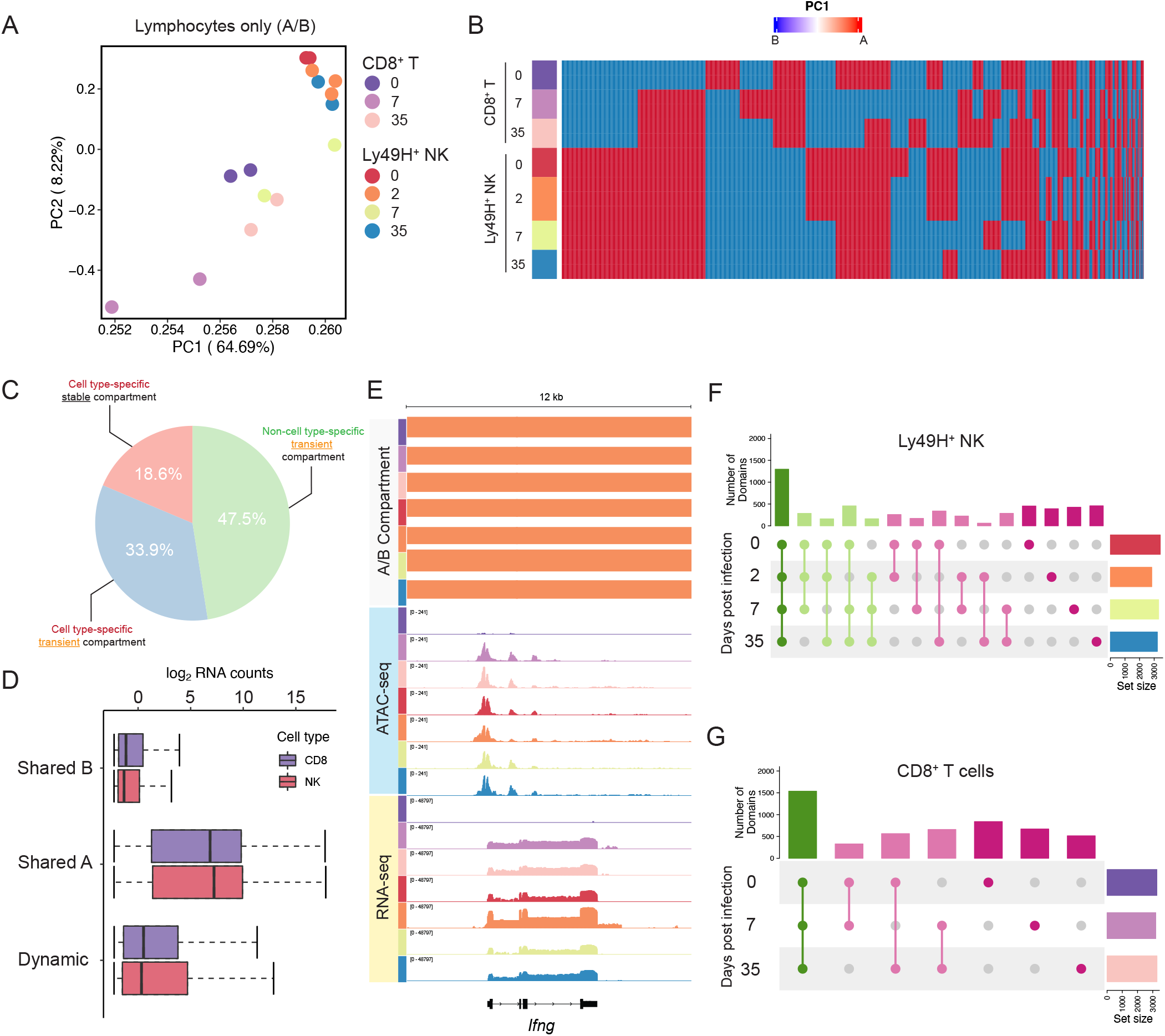
A/B compartments and cell type-specific TAD analysis. (A) Principal Component Analysis of A/B compartment profiles of Ly49H^+^ NK, CD8^+^ T across different time points after viral infection. (B) Heatmap of ‘dynamic’ compartments as binary values (A or B) ranked by the frequency of A/B compartment patterns. (C) Pie chart quantifying the percentage cell type-specific or non-cell type-specific compartments that are either stable (does not change across time points) or transient (flipped at least once during infection) within the ‘dynamic’ regions. (D) Average of log_2_ normalized RNA counts across different time points of genes that are located within ‘stable A’, ‘stable B’, or ‘dynamic’ compartments. (E) Genomic tracks depicting A/B compartment, chromatin accessibility (ATAC-seq) and transcript (RNA-seq) within the *Ifng* loci of CD8^+^ T cells (d0, 7, 35) and Ly49H^+^ NK cells (d0, 2, 7, 35) after infection. Orange bars represent A compartment, and light blue bars represent B compartment. (F) Upset plot of high-confidence TADs that are shared or distinct within Ly49H^+^ NK cells across different time points post MCMV. (G) Upset plot of high-confidence TADs that are shared or distinct within CD8^+^ T cells across different time points post MCMV.

**Supplementary Figure 5.**
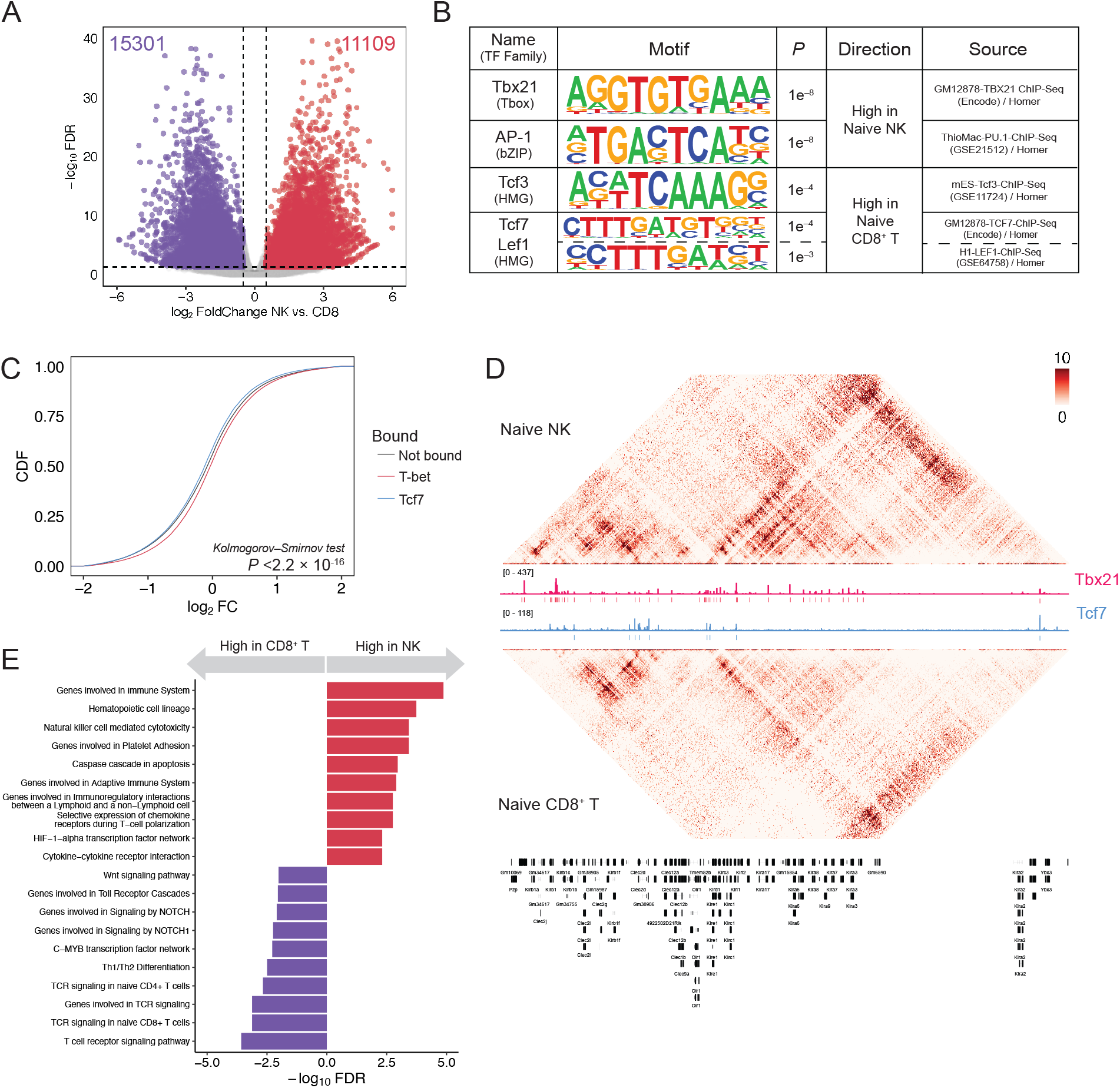
Distinct 3D chromatin landscape in naïve innate and adaptive lymphocytes. (A) Volcano plot on DI regions at 25kb comparing naïve CD8^+^ T and NK cells. Points highlighted are significant DI loops (p_adj_ < 0.05 & |log2FC| ≥ 0.5) that are either enriched in naïve CD8^+^ T cells (purple) or enriched in naïve NK cells (red). (B) Known motif analysis on DI regions between naïve CD8^+^ T and NK cells. (C) Cumulative distribution function (CDF) plot comparing DI regions that are either not bound or bound by TCF7 or T-BET. Two-sided Kolmogorov-Smirnov test was done on each pairwise comparison. In all comparison, *P* was found to be *<*2.2×10^−16^. (D) Contact heatmap matrix at 5kb resolution on naïve NK and CD8^+^ T cells, as well as T-BET ChIP-seq on naïve NK cells and TCF7 ChIP-seq on naïve CD8^+^ T cells. (E) Top 20 distinct pathway analysis of DI regions (p_adj_ < 0.05 & |log2FC| ≥ 0.5) between naïve CD8^+^ T and Ly49H^+^ NK cells as assessed by GREAT.

**Supplementary Figure 6.**
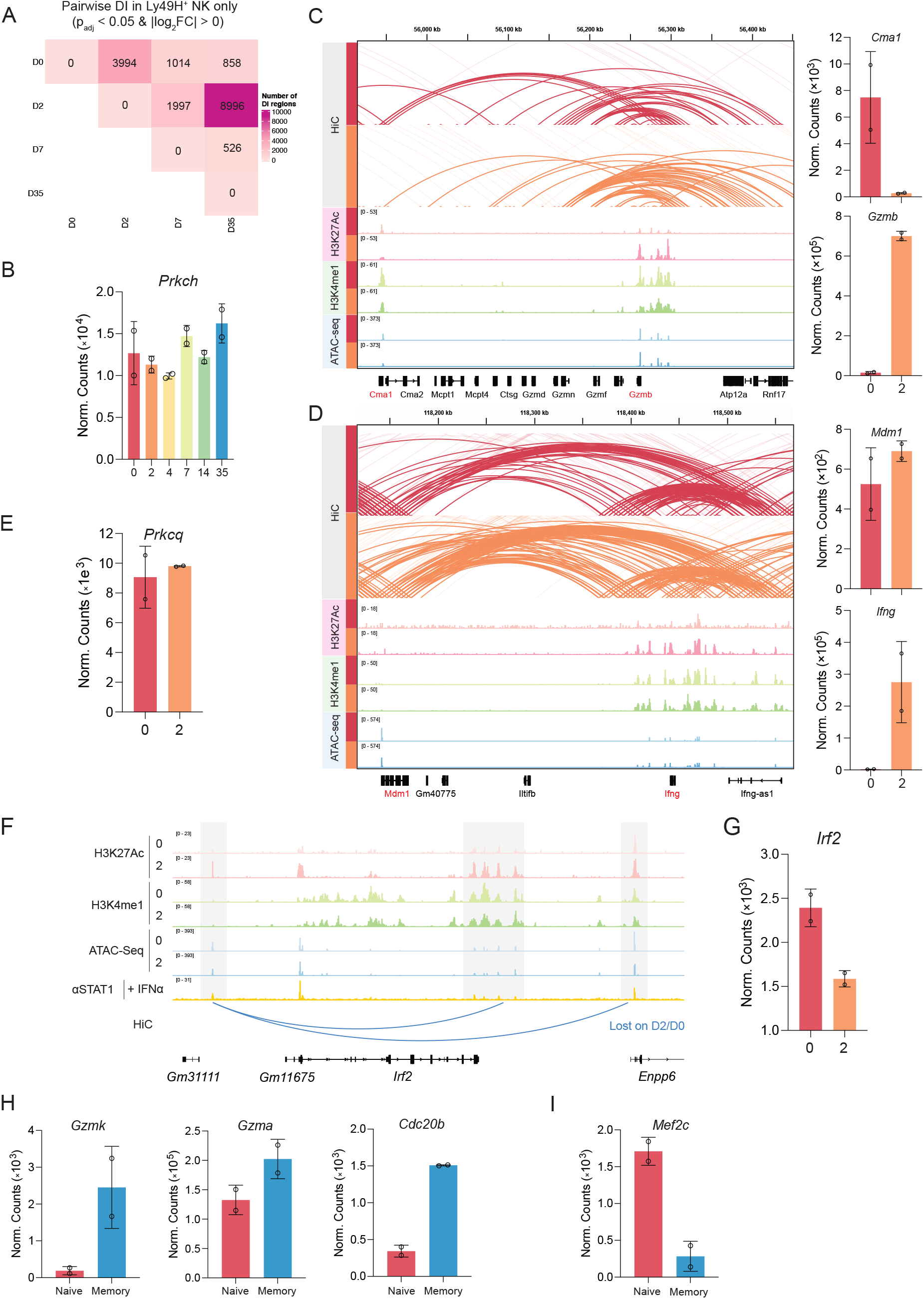
Three-dimensional chromatin dynamics in innate lymphocytes upon viral infection. (A) Pairwise comparison of DI regions in Ly49H^+^ NK only interaction atlas. (B) Normalized counts of *Prkch* in Ly49H^+^ NK cells across different time points post MCMV. (C) Genomic tracks depicting union of 5kb resolution significant HiC contacts, H3K27Ac, H3K4me1, ATAC-seq signals (day 0 and day 2 PI) of Ly49H^+^ NK cells around *Cma1* and *Gzmb* loci (left). Normalize RNA counts of *Cma1* and *Gzmb* on day 0 and day 2 PI in Ly49H^+^ NK cells. (D) Genomic tracks depicting union of 5kb resolution significant HiC contacts, H3K27Ac, H3K4me1, ATAC-seq signals (day 0 and day 2 PI) of Ly49H^+^ NK cells around *Mdm1* and *Ifng* loci (left). Normalize RNA counts of *Mdm1* and *Ifng* on day 0 and day 2 PI in Ly49H^+^ NK cells. (E) Normalized RNA counts of *Prkcq* on day 0 and day 2 PI in Ly49H^+^ NK cells. (F) Genomic tracks depicting H3K27Ac, H3K4me1, ATAC-seq signals (day 0 and day 2), and STAT1 ChIP-seq from *in vitro* stimulated NK cells, as well as gained DI regions on day 2 vs. day 0 in *Irf2* locus in Ly49H^+^ NK cells. (G) Normalized RNA counts of *Irf2* on day 0 and 2 PI in Ly49H^+^ NK cells. (H) Normalized counts of *Gzmk, Gzma, Cdc20b* transcripts in naïve and memory Ly49H^+^ NK cells. (I) Normalized counts of *Mef2c* transcripts in naïve and memory Ly49H^+^ NK cells.

**Supplementary Figure 7.**
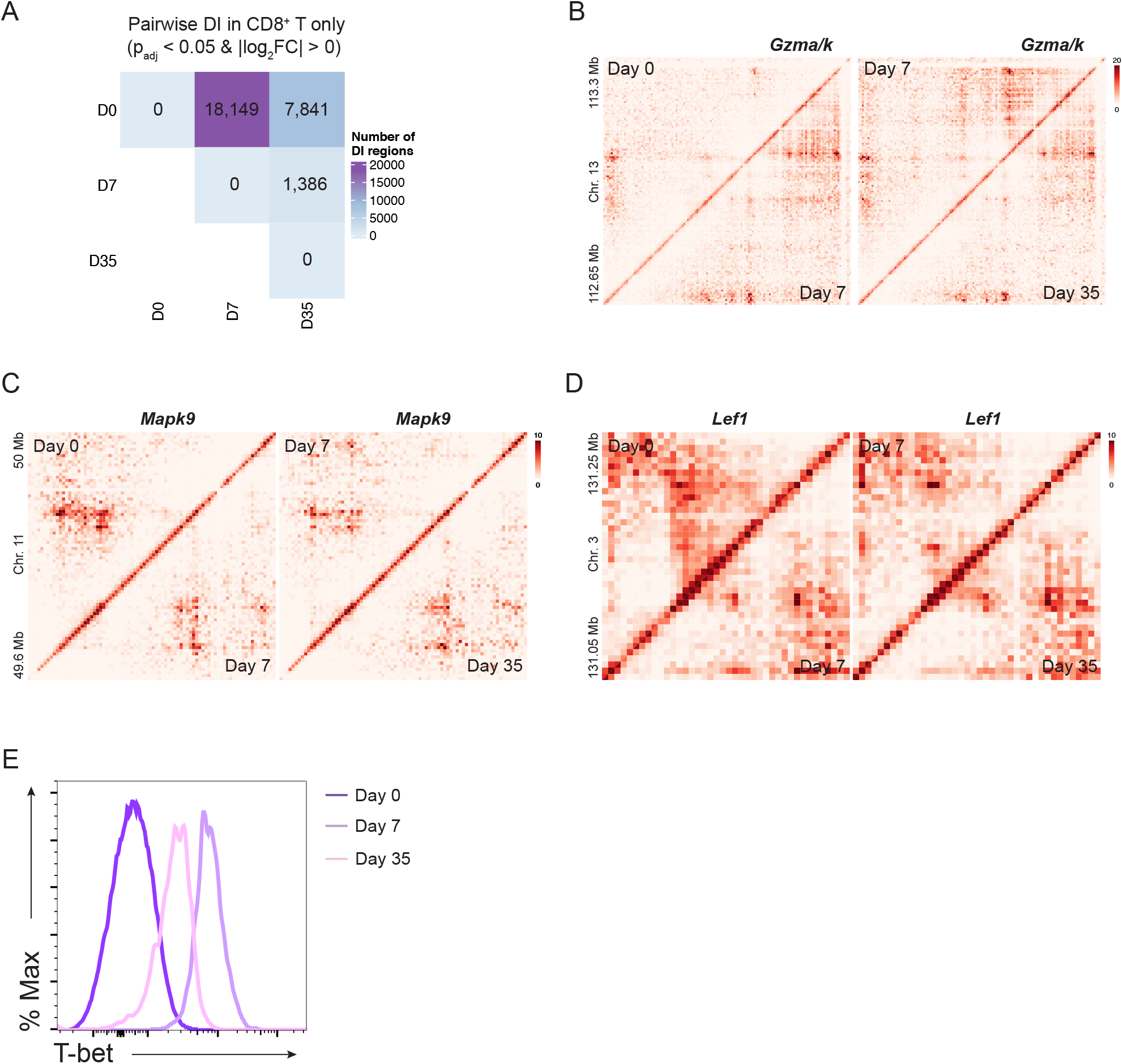
Three-dimensional chromatin dynamics in adaptive lymphocytes upon viral infection. (A) Pairwise comparison of DI regions in CD8^+^ T only interaction atlas. (B) Contract matrix heatmap near the *Gzma* and *Gzmk* loci in differentiating CD8^+^ T cells depicted as z-value. Gene label is located relative to the gene position in the genome. (C) Contract matrix heatmap near the *Mapk9* loci in differentiating CD8^+^ T cells depicted as z-value. Gene label is located relative to the gene position in the genome. (D) Contract matrix heatmap near the *Lef1* loci in differentiating CD8^+^ T cells depicted as z-value. Gene label is located relative to the gene position in the genome. (E) Histogram of T-BET expression as assessed by flow cytometry on MCMV-specific M45^H2-Db+^ CD8^+^ T cells at the indicated time points.

